# Dimensionality reduction and data integration for scRNA-seq data based on integrative hierarchical Poisson factorisation

**DOI:** 10.1101/2021.07.08.451664

**Authors:** Thomas Wong, Mauricio Barahona

## Abstract

Single-cell RNA sequencing (scRNA-seq) data sets consist of high-dimensional, sparse and noisy feature vectors, and pose a challenge for classic methods for dimensionality reduction. Such problems are compounded when dealing with composite data sets formed by different batches. We introduce Integrative Hierarchical Poisson Factorisation (IHPF), an extension of HPF that makes use of a noise ratio hyper-parameter to tune the variability attributed to batches *vs*. biological sources (cell phenotypes). We exemplify the application of IHPF under different data integration scenarios with varying alignments of batches and cell diversity, and show that IHPF produces latent factors that can be advantageously applied for cell clustering and visualisation. In addition, the extracted factors have a dual block structure in both cell and gene spaces with enhanced biological interpretability.

## 1. Introduction

Recent advances in sequencing technologies make it possible to profile biological processes at single-cell resolution, allowing the characterisation of the cell-to-cell variability that underpins numerous phenomena in biology and medicine. Single-cell RNA sequencing (scRNA-seq) has a wide range of applications measuring cell heterogeneity, from cancerous tumours to antibiotic resistance in bacterial populations [1, 2]. In scRNA-seq data sets, each sample is a cell described by a high-dimensional feature vector containing the transcription counts of tens of thousands of genes. Hence the *sample space* correspond to cells, and the *feature space* refers to genes. The feature vectors are discrete, noisy and sparse, with many zeros and low counts.

Due to the large number of features (i.e., genes), dimensionality reduction is a key ingredient in the analysis workflow of scRNA-seq data sets. Hence many computational pipelines are designed to extract a reduced set of informative and interpretable *latent factors* that capture the distinguishing characteristics of the cell types in the data set. However, the sparse and discrete nature of scRNA-seq data poses challenges to classic methods of dimensionality reduction, e.g., Principal Component Analysis (PCA) or Non-negative Matrix Factorisation (NMF). An additional challenge is posed by data integration. It is common to integrate data sets sequenced with different technologies and across different laboratories to validate hypotheses against previous findings, or to uncover rare biological signals that are difficult to observe in individual experiments. For example, researchers might combine a newly sequenced data set with existing reference data sets to identify cell types. Under such data integration scenarios, biologically meaningful signals must be separated from technical effects, such as differences in sequencing depth, quality control, or population effects (e.g., demographics of patients providing the samples) [3]. Such batch effects are often non-linear, and hence problematic for batch correction methods based on, e.g., PCA [4].

There are two major data integration tasks in single-cell genomics. One type of task is multi-omics integration, which aims to combine different modalities of data (e.g., transcriptomics and proteomics) through *a shared cell space*, as done by tools such as MOFA+ [5] and DIABLO [6]. A second integration task aims to combine data sets of the same modality but collected from different experiments, and correct batch differences *on a shared gene space*. Examples of these methods include scVI [7] and Scanorama [8]. Here, we focus on this latter task.

Several tools have been recently deployed for batch correction of scRNA-seq data ranging from traditional matrix factorisation, such as PCA or NMF, to deep learning methods, such as variational autoencoders. For a survey, see [4, 9]. A common characteristic of these methods is that they involve dimension reduction of the data followed by graph-based clustering using, e.g., k-nearest neighbour graphs. Batch correction is applied on the distance between cluster centroids under a suitable metric. However, these methods typically require long computational times, thus limiting their use for large-scale single-cell studies [10], and can lack robustness, with performance highly dependent on data pre-processing and gene filtering, on specific choices of hyper-parameters, and on fine-tuning for particular sequencing technologies [9].

An additional, desirable requirement in dimensionality reduction of scRNA-seq data sets is that latent factors are interpretable in both the cell and gene spaces, so that they can be used to generate biologically meaningful data-driven hypotheses linking gene expression and cell phenotypes. However, many of the current methods (e.g., PCA) lack interpretability due to the signed nature and lack of sparsity of the latent factor scores in the gene or cell spaces. Several methods have been proposed to improve interpretability. In particular, NMF ensures latent factors that learn a part-based structure, with non-negative scores, but with no guarantee of sparsity. Lack of interpretability is compounded, and especially relevant, when dealing with data integration of cell types across technical batches.

In this paper, we focus on data integration and dimensionality reduction of scRNA-seq data sets comprised of several batches, and consider this problem in the context of matrix factorisation methods that induce robust and interpretable latent factors to simplify downstream analyses.

To do so, we introduce Integrative Hierarchical Poisson Factorisation (IHPF), an extension to HPF that allows data integration and provides a robust, flexible and interpretable method for the analysis of multi-batch scRNA-seq data sets under different data integration scenarios.

The paper is organised as follows. Section 2 provides background on existing factorisation methods for dimensionality reduction pertaining to scRNA-seq analyses. Section 3 describes our algorithm for Integrative Hierarchical Poisson Factorisation (IHPF), the data sets, our numerical experiments, and we demonstrate how IHPF can be used to carry out dimensionality reduction under different data integration scenarios. In Section 4, we show that the latent factors obtained by IHPF provide a good basis for cell clustering and visualisation, and are also sparse and interpretable due to their dual block-structure in both cell and gene spaces. We conclude with a brief discussion in Section 5.

## 2. Background

### 2.1. Matrix factorisation and dimensionality reduction

Consider a data set consisting of *N* samples, each of them described by *M* ≫ 1 features. In the case of scRNA-seq, the *N* samples are cells of different types, and each of the *M* features is the transcription count of a particular gene. The data is compiled into a data matrix *X* of dimensions *N* × *M*.

Matrix factorisation methods attempt to find 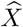, an approximation of the data matrix *X* in terms of two matrices *W* and *V* with reduced dimension (rank) *K ≪ M* such that their product is close to the original matrix:

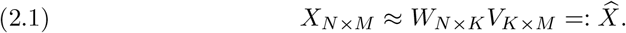

Here, the matrix *W* contains the cell scores and the matrix *V* contains the gene scores with respect to *K latent factors*. The latent factors thus recapitulate (approximately) the information contained in the *M* features, and each sample (cell) can be described by *K* coordinates.

PCA and NMF are two classic methods that produce such a matrix factorisation, differing in how the approximation (2.1) is computed. PCA minimises the Frobenius norm

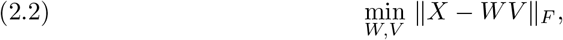

and it is well known that PCA approximations are obtained through the singular value decomposition (SVD) of the matrix *X*, i.e., the *K* factors are obtained from the top *K* singular vectors of *X* ordered in decreasing order of their singular values [11]. PCA-based methods have been widely used with scRNA-seq data sets, typically after applying a log-transformation to the raw data counts. However, the fact that the cell and gene scores (*W* and *V*) are signed and not sparse, together with the log-transformation, reduces the interpretability of the latent space.

The data matrix in scRNA-seq is non-negative element-wise (*X* ≥ 0); hence a factorisation that captures this property is desirable. NMF proposes such a factorisation: it minimises the Frobenius norm but imposing non-negativity (element-wise) of the factorisation matrices [12]:

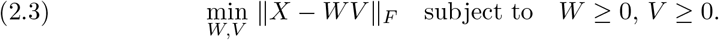

NMF learns part-based representations, which represent positive contributions of the latent factors to the samples [12], yet with no guarantee of sparsity.

### 2.2. Probabilistic viewpoint of factorization

Dimensionality reduction (and its associated matrix factorisations) also has probabilistic interpretations, specifically in terms of probabilistic factor analysis (FA) and hierarchical models.

From a probabilistic perspective, it has been shown [13] that PCA can be understood as a generative model where the samples are noisy observations of a combination of latent factors that are multivariate Gaussian [14, 15]:

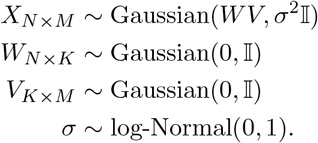

NMF also has a probabilistic interpretation related to Poisson factor analysis (PFA) [16]. It can be shown that under NMF, *X* is assumed to be a matrix of Poisson random variables with rates given by a factorisation into two matrices:

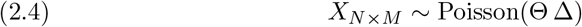

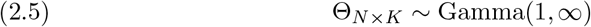

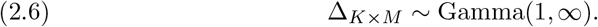

Here, Θ and Δ correspond, respectively, to cell and gene scores onto the *K* latent factors Θ and are Gamma distributed random variables with shape parameter 1 and infinite rate. Note that Poisson-Gamma random variables are related to the Negative Binomial distribution, which is widely used to model scRNA-seq counts.

Recently, this probabilistic interpretation inspired the Hierarchical Poisson Factorisation (HPF) [17], which uses a hierarchical Gamma prior:

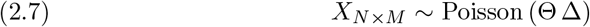

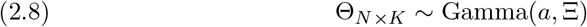

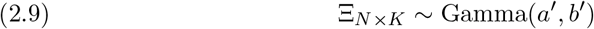

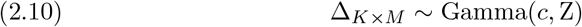

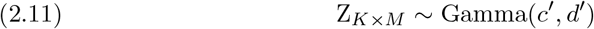

where the cell scores Θ follow a Gamma distribution with shape parameter *a* and a Gamma distributed rate Ξ. Similarly, the gene scores Δ are Gamma distributed with shape parameter *c* and a Gamma distributed rate Z.

HPF has been shown to perform better than NMF when applied to sparse data sets since the hierarchical Gamma prior is more flexible in modelling zero-inflated data counts [17].

### 2.3. Data integration

The problem of integrating data sets obtained with different technologies under diverse experimental settings poses additional challenges, as one is usually interested in capturing biologically meaningful signals in feature (gene) space under technical batch effects in sample (cell) space.

The classic matrix factorisation methods introduced in Section 2.1 have been extended for data integration. Traditionally, PCA has been used for batch correction in bulk RNA transcriptomics data, but it does not perform satisfactorily on scRNA-seq data due to the high sparsity of single cell data [2]. To circumvent these limitations, Scanorama [8] and BBKNN [18] apply graph-based algorithms on the principal components extracted by PCA to remove batch effects and improve alignment between cell types. BBKNN uses a k-nearest neighbour algorithm to build a graph to capture cell-to-cell similarity, attempting to balance the contribution from each batch. Scanorama identifies mutual nearest neighbours between batches and then finds the optimal order of batch alignment for panorama stitching, followed by a non-linear transformation to create the integrated latent space and batch-corrected gene expression matrices.

Integrative NMF (INMF) extennds NMF to harmonise data sets that share their gene space [19]. For *S* batches with non-negative data matrices 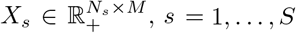, with 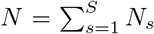, INMF finds a description in terms of K latent factors by obtaining non-negative matrices 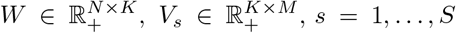, and 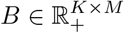 containing, respectively, cell scores, gene scores specific to each data set, and gene scores common to all data sets. The INMF factorisation minimises the loss function:

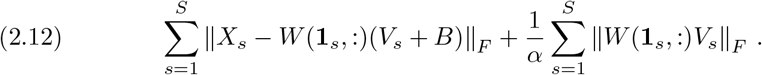

where **1**_*s*_ is the indicator function for cells in batch *s* and 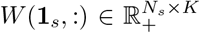 is the submatrix containing the cell scores for batch *s*. The hyper-parameter *α*, denoted as the *noise ratio*, regularises the magnitude of the components of the batches and assigns the variability in the data between cell types and batches.

Recently, deep learning has been used to perform dimensionality reduction and batch correction at the same time. scVI [7] uses a Variational Auto-Encoder (VAE) to learn a latent space representation of cells, modelling both library size and batch effects using a zero-inflated negative binomial distribution for gene expression counts. A significant drawback is that the latent space representation is not interpretable.

## 3. Methods

### 3.1. Integrative Hierarchical Poisson Factorisation (IHPF)

It has been shown that the performance of HPF can suffer when integrating data sets of different provenance (e.g., batches or other sample effects); hence various *ad hoc* methods (e.g., clustering of biologically related latent factors) have been applied to ameliorate batch effects [20].

Here we introduce an algorithm for *Integrative Hierarchical Poisson Factorisation* (IHPF), which extends the applicability of HPF to allow for systematic data integration, as follows.

Given data matrices 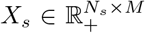, with *s* = 1,…, *S* indexing *S* batches, IHPF assumes a hierarchical Gamma prior on the cell scores Θ and it decomposes the gene scores into components Δ^(*s*)^, accounting for batch-specific effects, and a component B, accounting for shared gene effects. Both Δ^(*s*)^ and B are also described by a hierarchical Gamma prior. Then the IHPF model is given by:

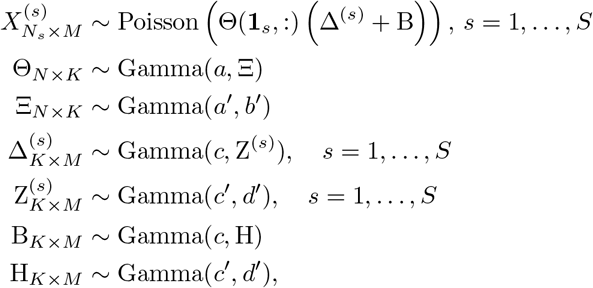

where, again, 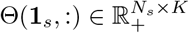 is the submatrix containing the cell scores of cells in batch *s*. The pseudo-code for the IHPF generative model is presented in Algorithm 3.1 and the IHPF algorithm is summarised in Figure 1.

**Fig. 1.**
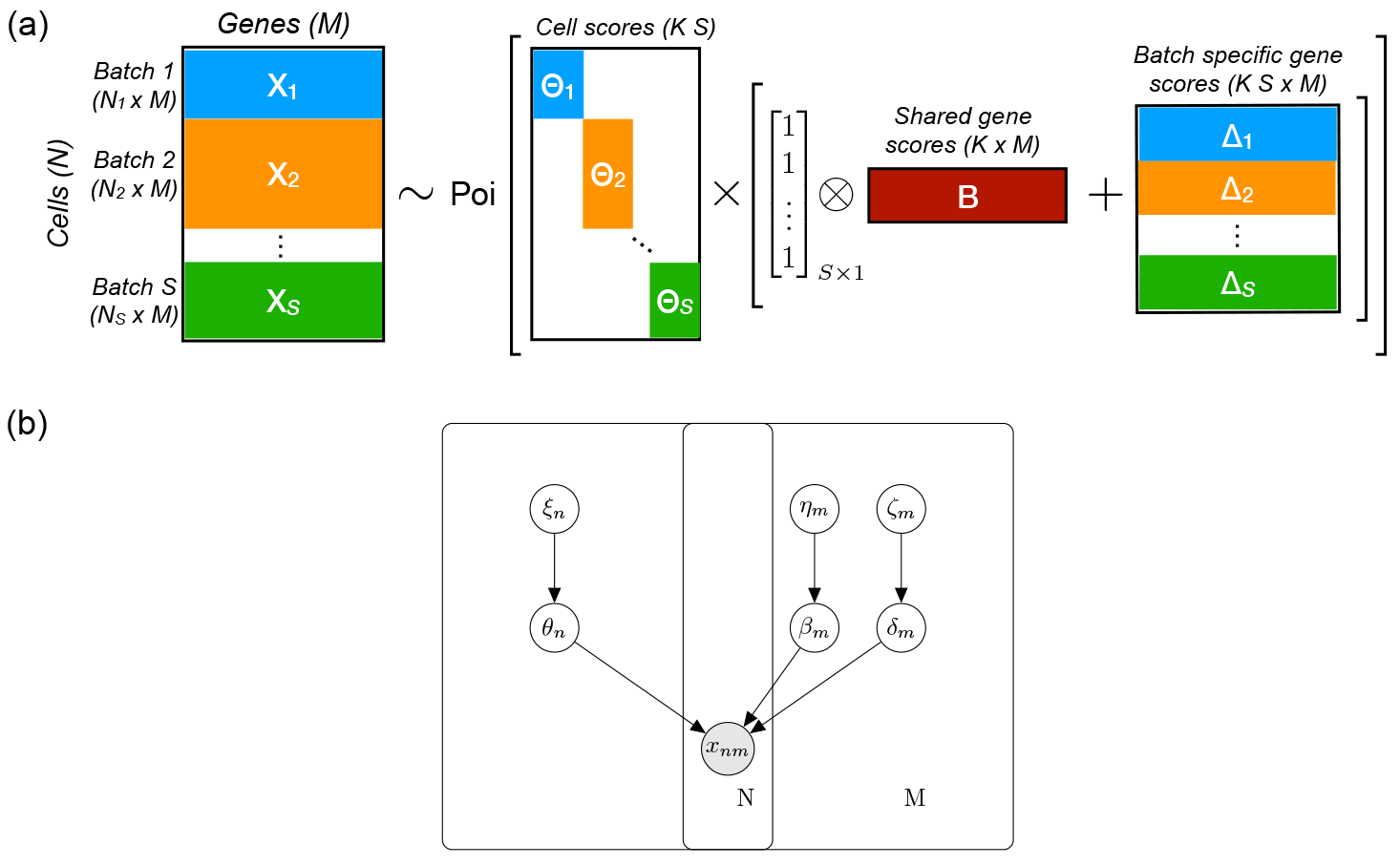
Two representations of the IHPF model. (a) IHPF can be viewed as a probabilistic matrix factorisation where the count matrix X, formed by the X_i_ of the S batches, is modelled as a Poisson distribution with weights given by a matrix factorisation with cell scores Θ_**i**_ together with batch-specific gene scores Δ_**i**_ and common gene scores B; (b) Graphical representation of the hierarchical IHPF model with N cells and M genes.

#### Algorithm 3.1

IHPF model for single-cell data.

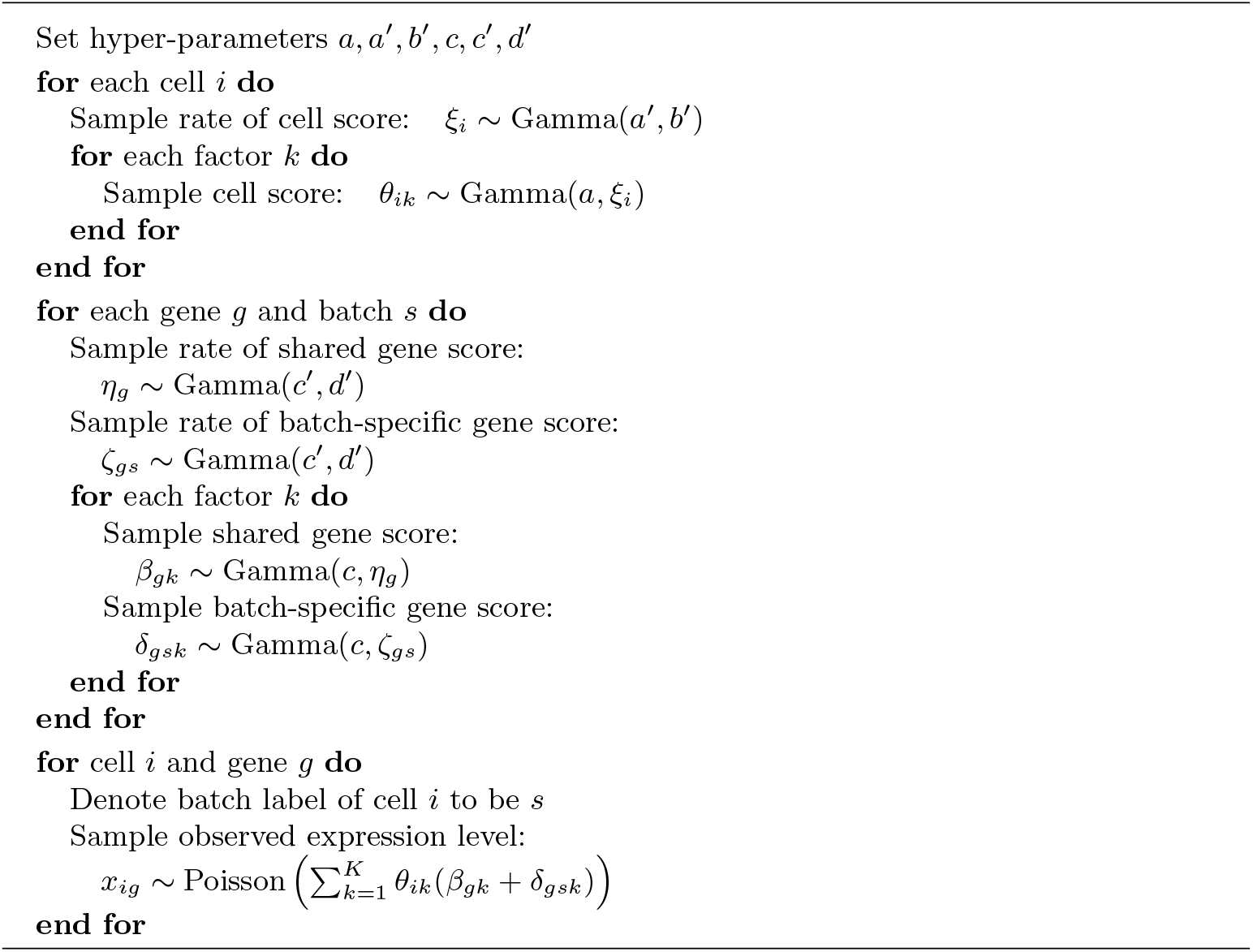

We use mean-field variational inference [21] to infer the IHPF model. Algorithm 3.2 describes the coordinate ascent algorithm [17] used to infer the shape and rate parameters of the six Gamma posteriors for gene and cell scores in Algorithm 3.1:

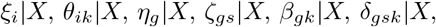

where each is Gamma distributed and the shape and rate parameters are defined accordingly for each posterior, e.g.,

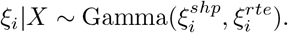

#### Algorithm 3.2

Coordinate Ascent Algorithm for likelihood inference in IHPF.

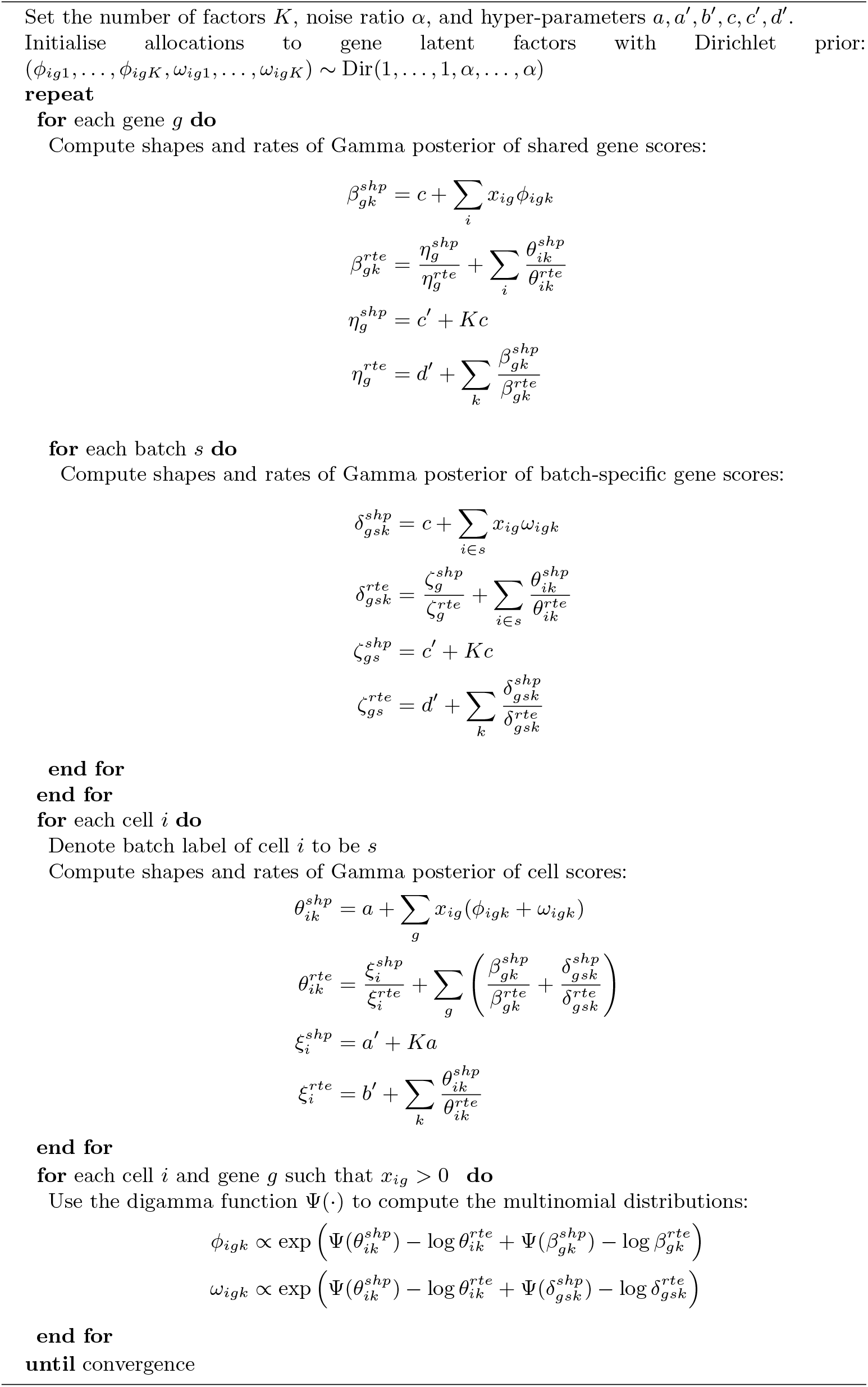

The multinomial distributions used in Algorithm 3.2 for the allocation of gene and cell counts to each latent factor are initialised with a Dirichlet prior with weights given by the noise ratio *α* ∈ [0, 1]. This hyper-parameter represents the relative importance of shared and batch specific gene latent factors, i.e., how much variance in the data is to be attributed to technical noise between data sets. Taking the limit *α* → 0 recovers the original HPF.

### 3.2. Data sets and data integration scenarios

We have collated from the published literature 16 scRNA-seq data sets of varying sizes and scope, and collected with different sequencing technologies (Table 1). Quality control is applied on the data sets by keeping only cells with at least 600 transcripts.

**Table 1.** Data sets used to benchmark various methods for data integration. Data sets are combined to produce integrated data sets to conform a specific scenario reflecting different assumptions on the biological relationship between data sets.

| ID | Description | Technology | Cells ($N$) | Genes ($M$) | Sequencing Depth | Scenario | Reference |
| --- | --- | --- | --- | --- | --- | --- | --- |
| 1 | Mouse Jurkat cells | 10X Genomics | 2885 | 32378 | 3060 | A | Zheng 1 [22] |
| 2 | Mouse 293t cells | 10X Genomics | 3257 | 32378 | 3168 | A | Zheng 2 [22] |
| 3 | Mouse 50% Jurkat/50% 293t cells | 10X Genomics | 3388 | 32378 | 3249 | A | Zheng 3 [22] |
| 4 | Human pancreas | InDrop | 8569 | 20126 | 2622 | B | Baron [23] |
| 5 | Human pancreas | Cel-seq2 | 2449 | 19140 | 3908 | B | Muraro [24] |
| 6 | Human pancreas | Fluidigm C1 | 638 | 25463 | 3294 | B | Lawlor [25] |
| 7 | Human pancreas | Smart-seq2 | 2989 | 26179 | 3342 | B | Segerstolpe [26] |
| 8 | Human pancreas | Celseq | 1276 | 20148 | 3196 | B | Grün [27] |
| 9 | Human CD19+ B cells | 10X Genomics | 2261 | 32378 | 1598 | C | Zheng 5 [22] |
| 10 | Human CD14+ monocytes | 10X Genomics | 295 | 32378 | 1824 | C | Zheng 6 [22] |
| 11 | Human CD4+ helper T cells | 10X Genomics | 3713 | 32378 | 1581 | C | Zheng 7 [22] |
| 12 | Human CD56+ natural killer cells | 10X Genomics | 6657 | 32378 | 1714 | C | Zheng 8 [22] |
| 13 | Human CD8+ cytotoxic T cells | 10X Genomics | 3990 | 32378 | 1509 | C | Zheng 9 [22] |
| 14 | Human CD4+CD45RO+ memory T cells | 10X Genomics | 3628 | 32378 | 1543 | C | Zheng 10 [22] |
| 15 | Human CD4+CD25+ regulatory T cells | 10X Genomics | 3365 | 32378 | 1609 | C | Zheng 11 [22] |
| 16 | Human PBMC | 10X Genomics | 2293 | 32378 | 1743 | C | Zheng 4 [22] |

We group data sets in Table 1 to conform three biological scenarios:

- *Scenario A:* a collection of data sets sequenced with the same technology and containing different abundance of the same cell types; here marker genes can easily tell the difference between cell types.
- *Scenario B:* a collection of data sets sequenced with different technologies and containing several cell types;
- *Scenario C:* a collection of data sets sequenced with the same technology in which there is little overlap in cell types between batches.

Table 2 summarises the three data scenarios. In our case, Scenario A consists of 3 batches of mouse cells (ID 1-3) sequenced with the 10X Genomics technology and comprising two cell types (293t and jurkat) mixed in different proportions. Scenario B consists of 5 batches of human pancreas cells (ID 4-8) of 10 different types obtained with different sequencing technologies. Scenario C consists of 8 batches (ID 9-16) sequenced with the same technology (10X Genomics) and comprising 6 different cell types highly concentrated in particular batches.

**Table 2.** Scenarios for data integration constructed from individual data sets from Table 1.

|  | Scenario A<br>(ID 1-3) | Scenario B<br>(ID 4-8) | Scenario C<br>(ID 9-16) |
| --- | --- | --- | --- |
| Total number of cells, $N$ | 9530 | 15921 | 26202 |
| Number of common genes, $M$ | 32643 | 15369 | 32643 |
| Number of batches, $S$ | 3 | 5 | 8 |
| Number of cell types, $C$ | 2 | 10 | 6 |
| AMI cell types *vs.* batches | 0.498 | 0.032 | 0.842 |

These three scenarios reflect experimental situations with different alignment between cell types and batch labels.

In Scenario A, batches and cell types have some alignment and, importantly, there is good overlap of cell types across batches. In Scenario B, the situation is complex, with many batches and cell types and with differing overlap of cell types across batches. In Scenario C, batches and cell types are highly aligned with little overlap of cell types across batches. To measure the alignment between cell types and batches, we use the Adjusted Mutual Information (AMI), as shown in Table 2.

### 3.3. Numerical Experiments and Code

To study data integration, we merge the corresponding data sets (keeping the intersection of genes across data sets) according to Scenarios A-C. Pre-processing of the single-cell data sets is done using Scanpy [28] and Scanorama [8], with details described in Section A.1. Our computational implementation of IHPF builds upon the original Python code for HPF in Ref. [17], and we use default hyper-parameters set in that reference. The data sets for the three data integration scenarios and the code implementing IHPF is available at https://github.com/barahona-research-group/scIHPF.

### 3.4. Data integration of scRNA-seq data sets using IHPF

#### Robustness of IHPF to hyper-parameters

First, we check the robustness of IHPF under changes of two (of its six) hyper-parameters: the cell shape parameter *a* and the gene shape parameter *c*. The other hyper-parameters (*a*′, *b*′, *c*′, *d*′) are fixed to the values recommended in the scHPF package [17]. The IHPF noise ratio *α* is set to 0.1.

We run IHPF for the three data integration scenarios varying the hyper-parameters *a* and *c* between [0.1, 1.1]. In all cases, we extract *K* = *C* factors, where *C* is the number of cell types present in the data. To evaluate the performance of model, we compute the negative mean log-likelihood of counts, under the Poisson assumption. Throughout this paper, we will always be dealing with Poisson log-likelihoods. Figure 2 shows that the performance of IHPF in all the data integration scenarios is highly robust to changes in the hyper-parameters *a* and *c*.

**Fig. 2.**
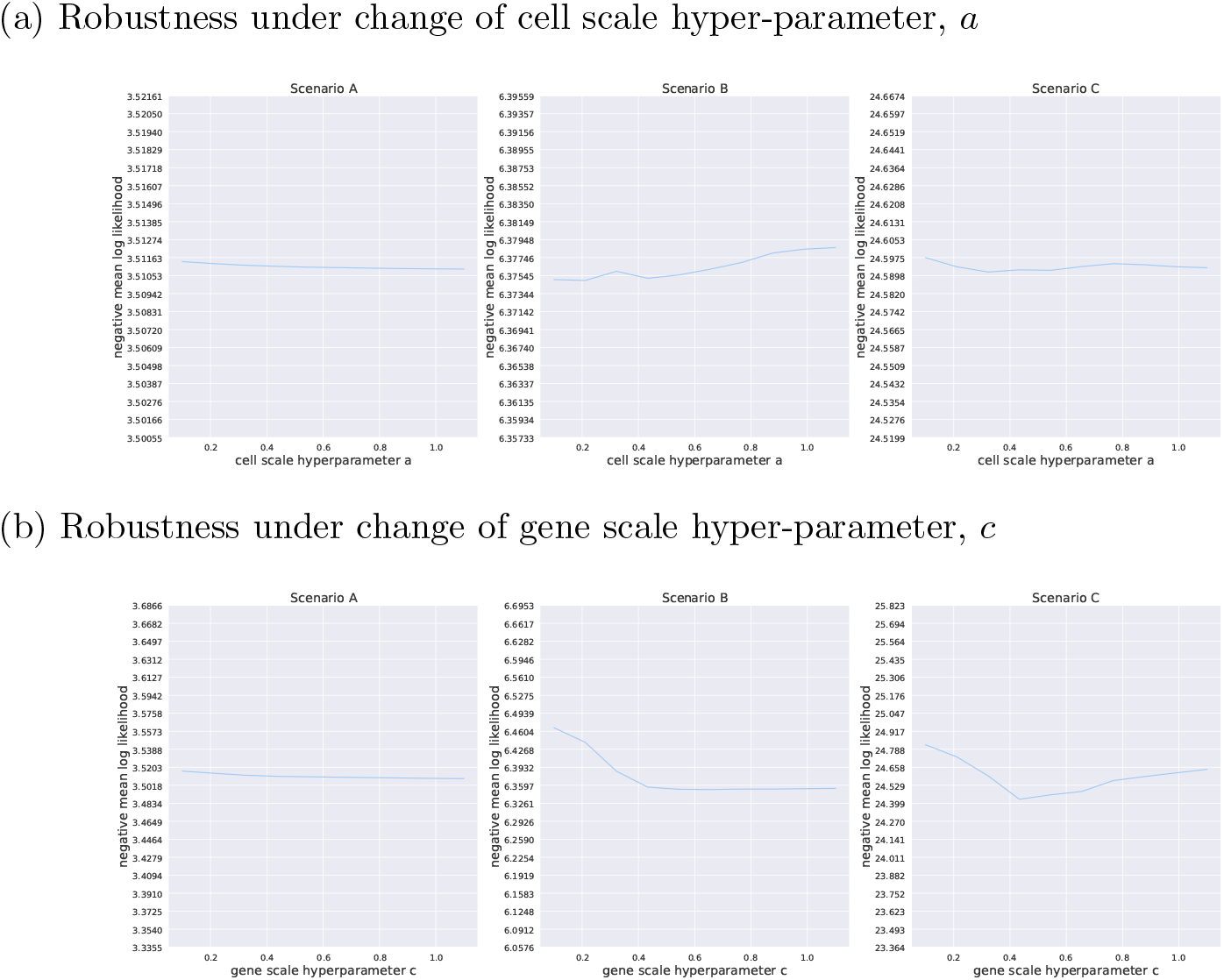
Negative Mean log-likelihood of IHPF for the three scenarios with varying hyperparameters: (a) cell scale hyper-parameter, a; (b) gene scale hyper-parameter, c. For all scenarios, varying the hyper-parameters has almost no effect on the model performance.

Once the robustness of IHPF to the hyper-parameters is established, we set *a* = *c* = 0.3 as suggested in the scHPF package [17] and keep these parameters fixed throughout the rest of the paper.

#### Selecting the IHPF noise ratio

The IHPF noise ratio *α* (see Algorithm 3.2) encodes how much of the observed data variance is attributed to the shared biological signal (i.e., cell types) or to batch-specific differences. When the noise ratio becomes zero, the batch labels are ignored and IHPF recovers the corresponding non-integrative result (i.e., HPF).

The noise ratio needs to be tuned for the specific data integration scenarios: a noise ratio that is too large will result in over-correction, so that the cell scores will not capture the biological difference between cell types; on the other hand, a noise ratio that is too small will result in extra cell clusters whereby cells with similar biology could be separated into smaller groups by batch.

While prior knowledge can be used to select the noise ratio (e.g., experimental design or specific biological hypotheses for an experimental scenario), we propose a statistical procedure that can be used to guide the choice of noise ratio. To do so, we employ a *leave-one-batch-out* cross-validation procedure, as follows. For each scenario and noise ratio, we apply IHPF on all but one of the batches. A new HPF model is trained using the learn shared gene space as initialisation, and we then project the leave-out batch on the shared gene space, and compute the reconstruction loss (negative mean Poisson likelihood) for that batch. We then repeat this procedure for each batch and compute the weighted average of the likelihood. The results are shown in Figure 3. (See Section A.2 for the results of all batches before averaging.) We then use the elbow method to identify the optimal noise ratio for each scenario, by taking the largest noise ratio until we experience a steep increase in reconstruction loss.

**Fig. 3.**
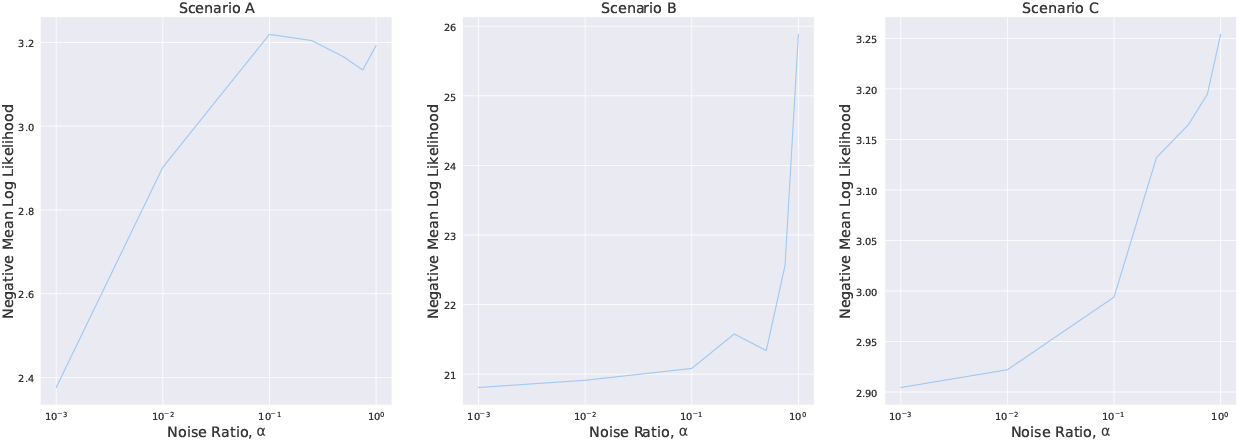
Cross-validation leave-one-batch-out likelihood for each data scenario, computed as the weighted mean of likelihoods for each batch, weighted by batch size. For every batch in each scenario, the IHPF model is fitted using the rest of the batches. This is then repeated for different noise ratios.

For Scenario A, there is a sharp increase of the negative likelihood even for small values of the noise ratio, hence there is no observed improvement with data integration and the optimal noise ratio selected is 0.001. For Scenario B, the noise ratio can be increased up to values of around 0.5 before the likelihood sharply deteriorates; hence the optimal noise ratio selected in this case is 0.5. For Scenario C, the optimal noise ratio selected is 0.01; hence in this case only mild data integration across batches is beneficial. Henceforth in the paper, we will fix these noise ratios for Scenarios A, B and C, unless otherwise indicated.

The optimal noise ratios selected are broadly consistent with our prior biological knowledge of the data sets. Indeed, batches in Scenario A similar to each other, batches in Scenario C are highly aligned with the cell types, and batches in Scenario B are both technically distinct and containing cell mixtures; hence data integration is more important in this last case. The noise ratio thus serves as an indication of the amount of data integration necessary for the particular collection of data sets, with the aim of taking into account batch variability yet still capturing the biological variability in the data.

## 4. Results

In this section, we first compare the quality of the latent factors obtained with different methods for the purpose of clustering cell types and visualisation, and consider the effect of the noise ratio for data scenarios with different alignment between cell types and batches. Secondly, we show that IHPF extracts sparse latent factors with a dual block structure with respect to both gene and cell spaces, thus enhancing their interpretability.

### 4.1. Using IHPF latent factors for cell clustering and visualization

Dimensionality reduction is a key step in computational pipelines for (unsupervised) clustering of scRNA-seq data sets. Ideally, one aims to extract a few latent factors that recapitulate the cell labels as fully as possible, and use the latent factors as data-driven coordinates on which to base classification, clustering, visualisations or coarse-grained descriptions that relate cell phenotypes to coherent groups of genes. PCA- or NMF-based factors have had only limited success with scRNA-seq data due to the high levels of noise and zeroes in the data [9].

To circumvent such limitations, complex pipelines have been developed including the use of similarity graphs and graph-based partitioning [8, 29, 30]. However, such methods encounter problems under data integration scenarios since the latent factors could capture mostly batch information, and in that case the clusters would fail to recapitulate the cell types. Rather than considering only complex multi-step computational pipelines with many parameters, we examine here if relatively simple data integration methods can extract latent factors that capture cell type variability in the presence of batch effects.

We evaluate five widely-used methods for dimensionality reduction and data integration based on different principles: Scanorama, BBKNN, scVI and INMF, along with our proposed method IHPF. The code for scVI [7], Scanorama [8] and BBKNN [18] were all downloaded from their corresponding Github pages, and we use recommended parameters and apply pre-processing steps on the count matrices following the recommendations by the authors. For INMF [19] we have coded a Python implementation compatible with scikit-learn, and set the noise ratio to the default value of 1.0, as suggested by the authors. For INMF and IHPF, we do not apply any pre-processing, other than the necessary quality control measures. Details of the pre-processing steps for each method can be found in the Supplementary Information.

#### Cell clustering from cell latent factors

We use each of the dimensionality reduction methods to extract *K* = *C* factors, where *C* is the number of cell types present in the data, and the row-normalised cell scores (e.g, from matrices *W* or Θ) are used to cluster the cells using k-means with *k* = *C*.

We first evaluate if the extracted cell latent factor scores induce well-separated clusters using the silhouette coefficient (SC), the (bounded) ratio of intra- and intercluster distances (–1 ≤ SC ≤ 1). For all methods other than BBKNN, SC is computed based on the cell latent factors (with latent dimension equal to the number of cell types in the scenario). For BBKNN, SC is computed based on the distance graph generated by the algorithm. Table 3 shows that, for all scenarios, IHPF gives the best separated clusters, as measured by the SC. On the other hand, methods that rely on Gaussian latent space and PCA, such as scVI and BBKNN give the least well-separated clusters, due to the mapping to non-sparse continuous variables. Scanorama and INMF provide an intermediate regime as a result of the graph-induced sparsity and the part-based property, respectively.

**Table 3.** Silhouette coefficient of the clusterings obtained by applying k-means clustering to K normalised cell latent factors obtained from five matrix factorisation models. Here, k = K = C.

| *Method* | Scenario A<br>(ID 1-3) | Scenario B<br>(ID 4-8) | Scenario C<br>(ID 9-16) |
| --- | --- | --- | --- |
| Scanorama | 0.447 | 0.292 | 0.335 |
| BBKNN | 0.007 | 0.079 | 0.094 |
| scVI | 0.107 | 0.185 | 0.075 |
| INMF | 0.458 | -0.229 | 0.446 |
| IHPF | 0.963 | 0.520 | 0.390 |

The SC computed above is an intrinsic measure of cluster separability, but does not evaluate the quality of the clusters with respect to the underlying cell types or, alternatively, to the batch labels. Therefore, to quantify the quality of the clusterings, we compute the Adjusted Mutual Information (AMI) between the cell clustering (i.e., k-means clusterings obtained from the cell scores) with respect to batch labels and cell types (Table 4).

**Table 4.**
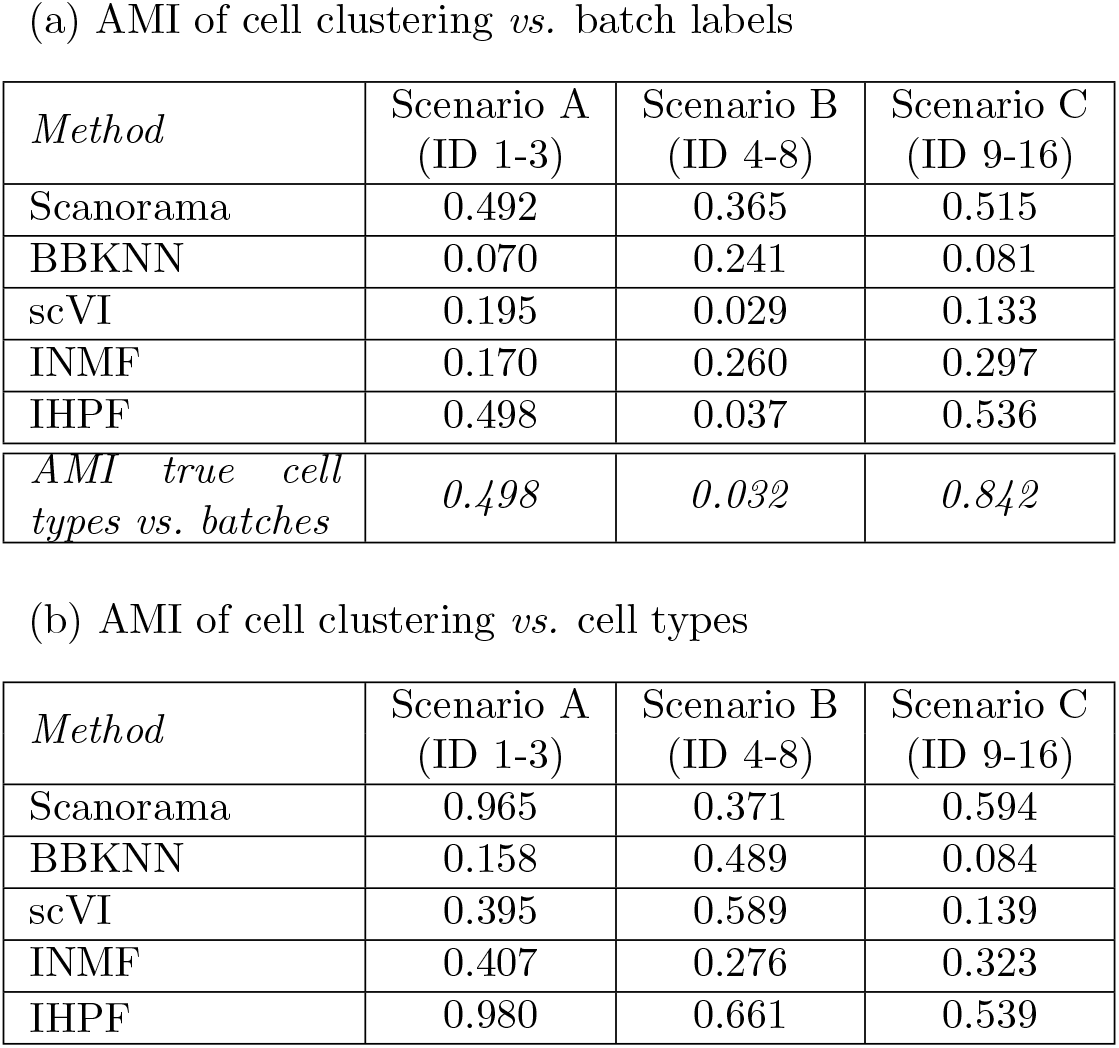
Adjusted Mutual Information (AMI) between the clustering obtained from latent factors versus (a) batches and (b) cell types. All the methods are applied without any gene selection. k-means is applied to K normalised cell latent factors obtained from each method, where the number of clusters k is set equal to the number of cell types C, i.e., K = k = C (see Table 2).

The AMI of the cell clusterings with respect to the batch labels (Table 4A) varies substantially for the different clustering methods and scenarios, but in general we observe that only IHPF is able to reduce batch effects in an adaptive manner, i.e., it only eliminates genuine batch effects that are not aligned with the cell types. This can be seen by comparing the AMI of IHPF clustering *vs*. batches to the AMI of true cell types *vs*. batches. The other methods either over-correct or under-correct batch effects, and fail to adapt to the underlying variation in the alignment of batches and cell types.

The objective of our clustering is to obtain a clustering from the latent factors that compares well with the cell labels (i.e., it is able to extract the biological signal in the presence of batch effects). The AMI of the cell clusterings *vs*. cell types (Table 4B) show that IHPF has the overall best performance. IHPF consistently outperforms INMF, BBKNN and scVI across all scenarios. Although their performance is comparable in Scenarios A and C, where integration is not crucial, IHPF outperforms Scanorama in Scenario B, where batch correction through adaptable data integration is beneficial. Therefore IHPF achieves good performance across different data integration scenarios with a simple model specification with a small number of parameters (especially compared to complex pipelines or deep learning methods such as Scanorama or scVI).

#### Visualisation based on cell latent factors

Another important use of latent factors is as part of visualisation pipelines used for biological exploration of data sets. Typically, the latent factors extracted with a dimensionality reduction method are used as inputs to nonlinear mapping algorithms (such as TSNE or UMAP) to obtain representations of the data on the plane. Such planar representations are broadly used to display high dimensional data and to guide biological hypotheses. The quality of the latent factors is thus an important factor to obtain useful planar representations.

As a brief illustration of the benefits of data integration, we compare the visualisations obtained with TSNE starting from PcA factors, as a simple baseline of dimensionality reduction without batch correction, compared to TSNE mappings using IHPF factors. Figure 4 shows the projections obtained from both IHPF and PCA factors using TSNE with default perplexity parameter [31]. Our results show that IHPF-based TNSE projections have better separated cell groups, which are less affected by potentially conflicting grouping by batches. This can be observed in the same projections coloured by batch label (Fig. 10 in SI), where we observe how the IHPF-based TSNE projections induce spatial groupings where cells of the same type but of different batches lie in close proximity, in contrast with the PCA groupings that are highly concomitant with batch labels. To see the same visualisations coloured according to batch type, see Section A.3 in the Supplementary Information.

**Fig. 4.**
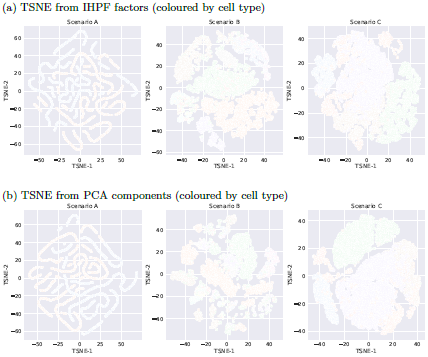
TSNE projections of Scenarios A-C obtained from latent factors from (a) IHPF and (b) PCA, and coloured by cell type.

### 4.2. IHPF induces block-structured factors in both gene and cell spaces

In biological applications, it is often desirable that the latent factors are both descriptive and interpretable. Ideally, a few, sparse factors should be able to recapitulate the original data. In this context, a sparse factor is one that has high scores concentrated in a few variables and low (or zero) scores for the remaining variables; hence the score matrices would have a well defined *block structure*. Such sparse factors can be the basis for clusters with improved biological interpretability, because they can link cell types to a reduced set of characteristic genes.

Most dimensionality reduction methods (e.g., PCA) are not designed to induce sparsity in the latent factors. On the other hand, IHPF is characterised by an intrinsic sparsity that makes its factors good candidates for improved interpretability. To explore this idea in more detail, we examine the block structure in the matrices of both cell and gene latent factor scores obtained with different methods.

#### Block-structure in cell space

Figure 5 shows the *cell scores* for Scenarios A-C obtained with PCA, INMF and IHPF. The block-structure present in all these matrices is a reflection of the fact that the cell latent factors capture information about cell types; indeed, these cell score matrices were the starting point for the clustering and visualisation results presented in Section 4.1.

**Fig. 5.**
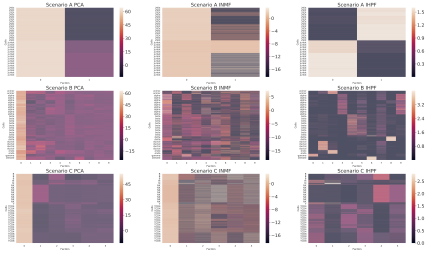
Cell latent factor scores (N × K matrices) obtained from PCA, INMF and IHPF for Scenarios A-C. The rows of the matrices correspond to cells sorted by cell type and the columns correspond to the latent factors. For each scenario, the number of latent factors is equal to the number of cell types (K = C). The block structure of the matrices is well aligned with the cell types, especially for IHPF.

Indeed, the silhouette coefficients (SC) for the k-means clusterings obtained from the cell scores of these three methods (Figure 6) confirm that IHPF cell scores lead to well separated clusters in all Scenarios, and PCA and INMF can also produce reasonable separability (especially for Scenario C, where data integration is not crucial).

**Fig. 6.**
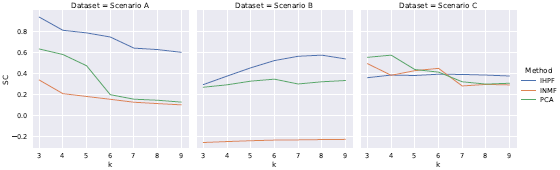
Silhouette coefficient (SC) of k-means clusterings (as a function of k) obtained from the N × K cell scores of PCA, INMF and IHPF for Scenarios A-C. Well separated clusterings, indicated by high SC values, are obtained from IHPF cell scores in all Scenarios.

#### Block-structure in gene space

The results above examine the structure of the latent factor scores in cell space, but in many cases it is advantageous to induce clusterings that also display high concentration and explainability in gene space. We have thus analysed the k-means clusterings obtained from the *K* × *M* matrices of gene scores (e.g., the matrix of common gene scores *B* in the case of IHPF). In contrast with the situation observed for the cell scores in Figure 5, we hnd that only the IHPF *gene scores* exhibit a well defined block structure across all three data scenarios (Figure 7). Consequently, the SC in Figure 8 shows that only IHPF gene latent factors induce well-separated gene clusters associated with the extracted latent factors. In summary, IHPF learns latent factors that have a dual block-structure, in both cell and gene spaces with the potential for enhanced explainability and biological interpretability by linking cell types to gene clusters, as explored below.

**Fig. 7.**
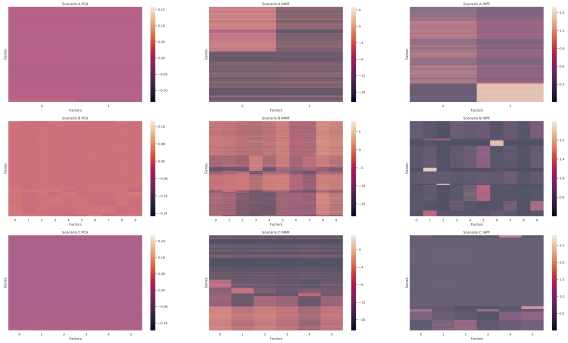
Gene latent factor scores (K × M matrices) obtained from PCA, INMF and IHPF for Scenarios A-C. As above, K = C. The rows (genes) are ordered using spectral co-clustering for ease of visualisation. There is only well-aligned block structure for the IHPF gene scores.

**Fig. 8.**
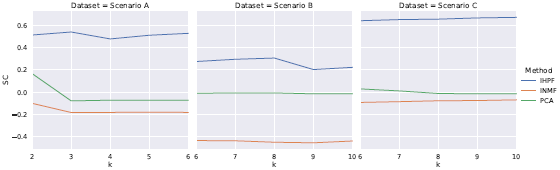
Silhouette coefficient (SC) of k-means clusterings (as a function of k) obtained from the gene scores of PCA, INMF and IHPF for Scenarios A-C. Only IHPF gene factors lead to well separated clusters with high SC values whereas INMF and PCA gene factors lead to poor clusters with negative values of SC.

#### Biological homogeneity of extracted gene clusters

To characterise further the biological interpretability of the gene clusters extracted with IHPF, we proceed as follows. We start by selecting the number of clusters that gives the highest Silhouette coefficient (SC) of the gene scores (Figure 8). For Scenario A, the optimal number of gene clusters is 3. For Scenario B, the optimal number of gene clusters is 8. For Scenario C, the optimal number of gene clusters is 10. Note that the number of gene clusters is different from the number of cell types, as there is no one-to-one mapping between cell types and gene groups—a cell type can be linked to multiple gene groups and *vice versa*.

We then compute the biological homogeneity index (BHI) [32, 33] of the obtained gene clusterings. The BHI is a measure of within-cluster biological similarity of the Gene Ontology (GO) terms associated with the genes present in each cluster. Specifically, for a clustering 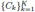 with *K* clusters *C*_*k*_, the BHI is given by:

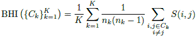

where *n*_*k*_ is the size of cluster *C*_*k*_, and *S*(*i, j*) is the biological similarity between gene *i* and gene *j*, obtained using an information-theoretic semantic similarity based on GO terms. The BHI gives a score between 0 and 1 [34]. For further details on the calculation of the BHI, see Section A.4 in the Supplementary Information.

For each gene clustering, we also compute the BHI of 100 randomised surrogate clusterings with the same number and sizes of clusters. We then compute the z-score and p-value of the BHI of the IHPF clustering against the ensemble of random clusterings to obtain a level of significance for the obtained clustering [35].

The results in Table 5 show that the clusterings obtained from the IHPF gene scores for Scenarios B and C are significant, with small p-values. Hence the gene clusterings correspond to meaningful biological groupings of genes.

**Table 5.** The z-scores and p-values for the BHI of the gene clusterings obtained from IHPF gene factors. The gene clusterings were obtained using k-means with k = 3,8, 10 for Scenarios A, B and C, respectively.

|  | Scenario A | Scenario B | Scenario C |
| --- | --- | --- | --- |
| z-score | 1.982 | 2.506 | 4.356 |
| p-value | 0.024 | 0.0061 | 0.0000066 |

## 5. Discussion

Here we have presented a computational method to carry out dimensionality reduction and data integration for single-cell RNAseq data based on integrative hierarchical Poisson factorisation (IHPF). The proposed algorithm extends HPF and performs data integration by modelling the technical noise between data sets with the use of latent factors adapted to batch labels.

As in other data integration methods (e.g., INMF), the noise ratio hyper-parameter can be tuned to accommodate the amount of variability ascribed to batch effects *vs*. biological effects (i.e., cell types). When the noise ratio becomes negligible, the batch labels are ignored and IHPF recovers the results of HPF. The noise ratio can be adjusted depending on the different experimental data integration scenarios. In cases when there is good overlap of cell types across batches and a large set of common genes (Scenario A), or the other extreme when batches and cell types are highly aligned with little overlap of cell types across batches (Scenario C), data integration brings little advantage. In that case, IHPF indicates that a close to zero noise ratio is appropriate, and still provides very good performance. On the other hand, in more complex situations (where batches are technically different and contain a varied mixture of cells), IHPF offers a flexible and controlled way to correct the effect of batch effects and can outperform other methods. We note that IHPF has no requirements on library sizes of the cells in the data sets, unlike PCA-based methods such as Scanorama, which requires pre-processing of cells to have normalised sizes. In addition, IHPF is shown to recover factors with dual block structure in both the cell and gene spaces, thus enhancing the explainability of latent factors in terms of groups of genes with biological content. We have made IHPF available as a Python package for use by the community.

There are several directions for extensions and improvement of this work. Currently, IHPF can only correct differences between data sets that are categorical in nature, such as batch effects. When the differences are continuous factors, such as population effects, a different model is needed. Further, the latent factors obtained with IHPF could also serve as the seed for more sophisticated classification and clustering methods, including variational autoencoders or graph-based multi-scale clustering methods [30, 36, 37].

## Acknowledgments

We thank Dr Zijing Liu for helpful discussions. We acknowledge funding by the Wellcome Trust under Grant 108908/B/15/Z, and by the EPSRC under grant EP/N014529/1 funding the EPSRC Centre for Mathematics of Precision Healthcare at Imperial.

## Appendix A. Supplementary Information

### A.1. Data pre-processing for the different scRNA-seq methods

Our pre-processing starts with quality control on the data sets, whereby we filter cells with a low transcript count. For each of the computational methods, we follow the preprocessing steps recommended by the original authors using Scanpy [28]. Specifically, we use the following functions:

- Scanorama: sc.pp.recipe_zheng17
- scVI: sc.pp.normalize_total, sc.pp.loglp
- BBKNN: sc.pp.recipe_zheng17
- INMF: None
- IHPF: None

### A.2. Leave-One-Batch-Out cross-validation

Detailed results of the leave-one-batch-out cross-validation likelihood for each of the batches are shown in Figure 9. The weighted average of these are presented in Figure 3 in the main text.

**Fig. 9.**
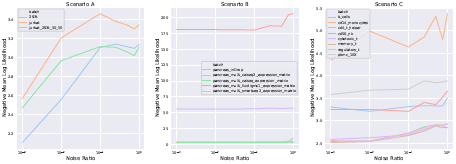
Cross-validation negative mean Poisson likelihood for each scenario. For every batch in each scenario, IHPF model is fitted using the rest of the batches for different noise ratios.

### A.3. Visualisation

Additional results on the visualisation of data sets using TSNE and PCA with cells colored by batch labels are shown in Figure 10.

**Fig. 10.**
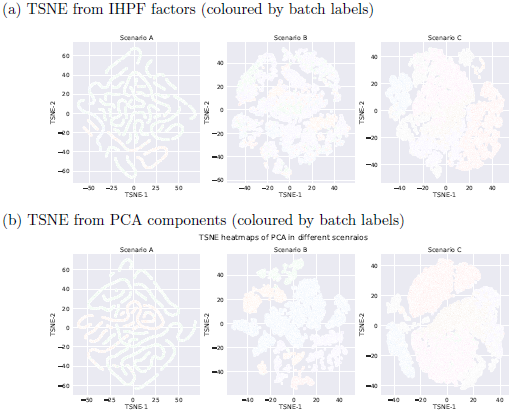
TSNE projections of Scenarios A-C obtained from latent factors from (a) IHPF and (b) PCA, and coloured by batch label.

### A.4. Computation of BHI

The genes in Scenarios A, B and C are mapped to GO terms using the org.Hs.eg.db annotation package. A total of 60.88 % of genes in Scenario A, 98.55 % of genes in Scenario B and 60.88 % of genes in Scenario C are mapped to a valid GO term. After that, the BHI is computed using the R package clValid [32].

